# Small molecule and cell contact-inducible systems for controlling expression and differentiation in stem cells

**DOI:** 10.1101/2024.03.12.584464

**Authors:** Sarah S. Soliman, Devan H. Shah, Hana El-Samad, Zara Y. Weinberg

**Affiliations:** Department of Biochemistry and Biophysics, University of California, San Francisco, CA; UC Berkeley-UCSF Graduate Program in Bioengineering, University of California, Berkeley, CA; Cell Design Institute, University of California, San Francisco, CA; Chan-Zuckerberg Biohub, San Francisco, CA; Altos Labs, Redwood City, CA

## Abstract

Synthetic developmental biology uses engineering approaches to understand multicellularity with goals ranging from recapitulating development to building synthetic organisms. Current approaches include engineering multicellular patterning, controlling differentiation, and implementing cooperative cellular behaviors in model systems. Synthetic biology tools enable these pursuits with genetic circuits that drive customized responses to arbitrary stimuli, synthetic receptors that enable orthogonal signaling channels, and light- or drug-inducible systems that enable precise spatial and temporal control of cell function. Mouse embryonic stem cells (mESCs) offer a well-studied and genetically tractable pluripotent chassis for pursuing synthetic development questions however, there is minimal characterization of existing synthetic biology tools in mESCs and we lack genetic toolkits for rapid iterative engineering of synthetic development workflows. Here, we began to address this challenge by characterizing small molecule and cell contact-inducible systems for gene expression in and differentiation of mESCs. We show that small molecule and cell-contact inducible systems work reliably and efficiently for controlling expression of arbitrary genetic payloads. Furthermore, we show that these systems can drive direct differentiation of mESCs into neurons. Each of these systems can readily be used on their own or in combination, opening many possibilities for studying developmental principles with high precision.

## Introduction

Synthetic developmental biology seeks to decipher the principles of developmental processes using a build-to-understand approach^1^. This entails building synthetic systems that recapitulate multicellular developmental paradigms such as differentiation, patterning, and cooperative cellular behaviors^2–7^. The field of synthetic developmental biology is still nascent, but holds immense promise spanning from controlling tissue development in a dish to engineering fully synthetic tissues and building synthetic organisms^8,9^.

Synthetic biology tools are essential for synthetic development and have been used in practically every landmark study in this field^4–6,10–13^. Synthetic biology has produced a repertoire of sophisticated tools for controlling cellular behavior that enables customized responses to arbitrary stimuli including synthetic receptors that enable orthogonal signaling channels and light- or drug-inducible systems that enable precise spatial and temporal control of cell function^14–20^. Cell engineering has been advanced through packaging these tools into robust modular toolkits such as the Mammalian Toolkit^21^. Adapting and evaluating these tools in developmental model systems will facilitate the goals of synthetic development.

When evaluating new tools in a synthetic development context, reprogramming is an accessible and well-studied testbed for applying synthetic biology tools^22^. Overexpression of transcription factors has been heavily used in the field of stem cell differentiation^23–26^. For example, multiple studies have demonstrated that the proneural transcription factor Neurogenin-2 (Ngn2) is necessary and sufficient to specify glutamatergic neuronal identity^27,28^. Efforts to control transcription factor overexpression in stem cells have primarily used the artificial transcriptional regulator TetR which is derived from microbial transcription factor and induced by the small-molecule antibiotic, doxycycline^29,30^. In recent examples, the synthetic receptor synNotch has been used in mESCs^31^ and human embryonic stem cells^32^ (hESCs) to drive neuronal differentiation.

The number of tools available to synthetically rebuild development are lacking in terms of the available breadth of small molecule and cell contact-inducible systems and the rapid iterability in which these systems can be incorporated into developmental models. To address this, we sought to expand the synthetic developmental toolbox by engineering synthetic biology tools into embryonic stem cells. While recent approaches in synthetic biology have elegantly recapitulated development in simple cellular systems, additional approaches are needed to fully realize the potential of synthetic developmental biology. Robust modular tools for engineering cellular behavior have had important impacts in yeast and mammalian cell engineering and we sought to bring those to stem cells. Here we describe an approach for rapid engineering of multiple inducible systems into mouse embryonic stem cells (mESCs). We show that these systems work reliably and efficiently in controlling expression of arbitrary genetic payloads. Furthermore, we show that some of these systems are capable of driving direct differentiation of mESCs into neuron-like cells, opening up questions about the quantitative requirements of these differentiation programs. The components and approaches we describe can readily be used on their own or in combination to engineer mESCs in synthetic development studies and further expand the toolkit for studying developmental principles with high precision.

## Results

### Orthogonal small molecule synthetic transcription factor systems induce BFP expression in mESCs

We first sought to build and test small molecule inducible synthetic transcription factor systems in mESCs. Inspired by the clinically-relevant synthetic zinc finger transcription regulators (synZiFTRs) system^18^, we adapted three different drug-inducible control switches for use in mESCs. The first of these drugs is 4-hydroxytamoxifen (4OHT), that selectively modulates the nuclear availability of molecules fused to a sensitized variant of the human estrogen receptor ERT2^33,34^. The second drug is abscisic acid (ABA), which mediates conditional binding of complementary protein fragments (ABI and PYL) from the ABA stress response pathway to reconstitute an active transcription factor^35^. The third drug is grazoprevir (GZV), a protease-inhibitor, that when added stabilizes transcription factors incorporating the hepatitis C NS3 self-cleaving protease domain^36,37^.

To ensure orthogonality of our transcriptional system from the murine genome, we employed the GAL4 upstream activating sequence (UAS) from yeast as our inducible promoter^38,39^. We selected a version of the GAL4 UAS that is coupled to the minimal promoter ybTATA^40^ which performs well in mammalian systems with low background activation, high signal to noise, and tunability^20,41,42^. The GAL4 UAS-ybTATA system has not been used in mESCs and therefore it would represent an expansion to the orthogonal transcriptional regulatory elements toolkit. Using the Mammalian Cloning Toolkit^21^, we constructed our synthetic transcription factor system using the GAL4 DNA binding and the transactivation domains of either VP16 (fused to ABI/PYL) or VP64 (4X tandem VP16, fused to ERT2 and NS3) coupled to a constitutively expressed mCherry as a marker of construct expression (Figure 1A, Supplemental Figure 1A)^43,44^.

**Figure 1:**
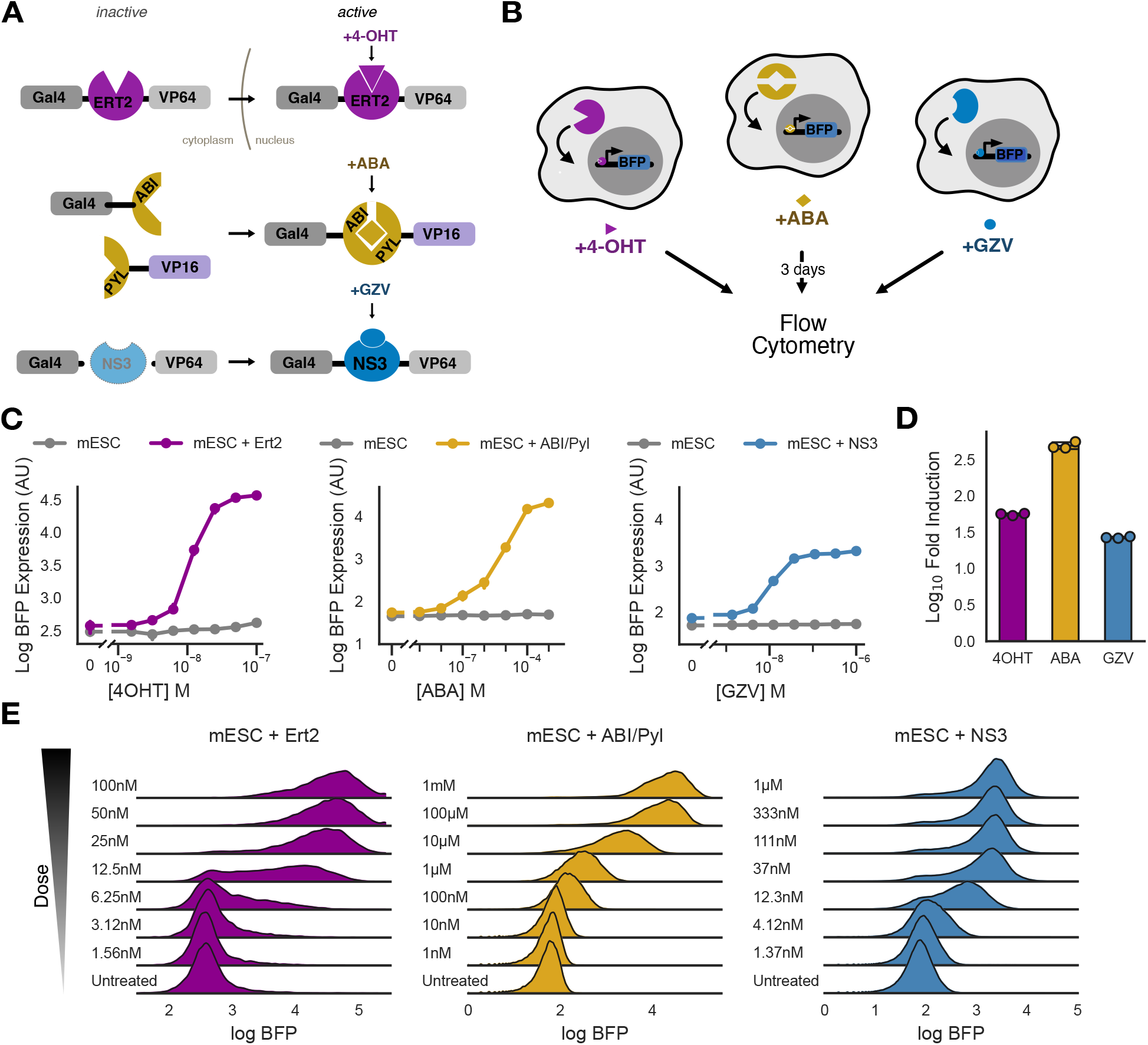
Orthogonal small molecule synthetic transcription factor systems induce BFP expression in mESCs. A) Schematic of small molecule synthetic transcription factor systems mechanism of action. B) Schematic of small molecule synthetic transcription factor systems assay. C) Dose response of curves of median log BFP expression for each small molecule synthetic transcription factor system. D) Median log_10_ fold induction levels for each system. E) Histograms describing population density estimates of each small molecule inducible system across doses.

To test the function of each of these systems, we used an output genetic cassette driving inducible TagBFP downstream of a 5X tandem GAL4 UAS sequence and a ybTATA minimal promoter alongside a constitutively expressed mCitrine^20,41^ (Supplemental Figure 1B). We used lentiviral transduction to introduce this cassette into mESCc and then sorted a mixed population of cells based on similar mCitrine expression. We then stably integrated our synthetic transcription factor systems into the output circuit mESC line by lentiviral transduction and sorted cells based on similar mCherry expression.

We cultured mESCs expressing each system with different concentrations of their specific inducer for three days, and assessed drug-dependent expression of TagBFP via flow cytometry (Figure 1B). The three inducible systems exhibited titratable control of TagBFP output, minimal leakage relative to TagBFP-only cells and strong dynamic ranges (Figure 1C). Each of these systems exhibited strong levels of induction (Figure 1D). These systems exhibited different dose-dependent population distribution dynamics within mESCs. Across a titratable dose, both ERT2 and NS3 shifted individual cells between discrete ON and OFF states while ABI/PYI showed a continuous response in individual cells (Figure 1E). These data indicate that these three orthogonal small molecule synthetic transcription factor systems work reliably and efficently to induce TagBFP expression in mESCs posing the potential to be used in more elaborate gene switches and in controlling more complex genetic payloads.

### Juxtracrine-inducible systems drive BFP expression in mESCs

Juxtacrine signaling is an important mechanism in development and physiology that allows cells to identify their tissue context and coordinate their behavior^45,46^. Recognition of cell surface antigens has been exploited in cell engineering to modulate pathways and drive different cellular behaviors^19,47,48^. SynNotch and SNIPR are designed juxtacrine signaling receptors that have proven useful in cell engineering applications. SynNotch-dependent induction has been used successfully to control cell patterning and drive differentiation in multiple cell types^31,32,49^. Although work in hESCs used lentivirus^32^, previous implementations of this system in mESCs required single site genomic integration and subsequent single cell sorting and clone selection to achieve functional engineered cells^31^. We sought to improve juxtacrine inducible signaling in mESCs by generating an implementation which could be prototyped rapidly and could produce high levels of its payload genes.

SynNotch and SNIPRs are both synthetic proteolytic receptors which drive transcription downstream of extracellular antigen recognition via customized sensing domains (Figure 2A)^19,20^. SNIPRs were developed by exploring the ability of a library of chimeric proteins to overcome the high background activity of synNotch in some cell types. We hypothesized that the functional differences between SNIPRs compared to the original synNotch might position SNIPRs as better suited for rapid iteration in engineering mESCs. We selected a SNIPR here that proved effective in recent studies in T cells^50^ containing the human CD8a hinge domain, human Notch1 transmembrane domain, and human Notch2 juxtamembrane domain.

**Figure 2:**
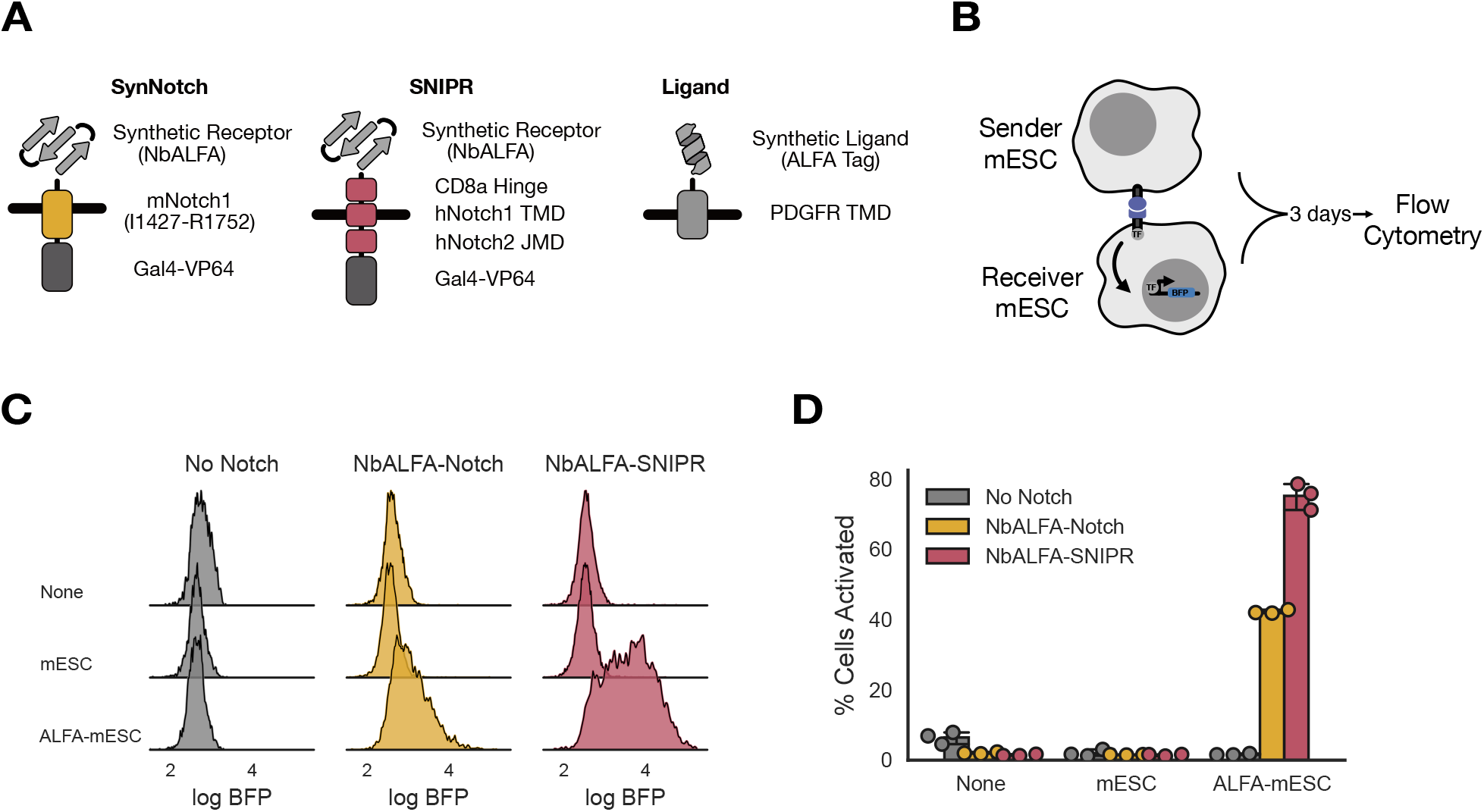
Juxtracrine inducible systems activate BFP expression in mESCs. A) Schematic of components of synNotch, SNIPR and ligand. B) Co-culture assay. C) Histograms for population density estimates for BFP expression comparing No Notch, NbALFA-Notch, and NbALFA-SNIPR when cultured alone (None), with wildtype mESCs, or ALFA-mESCs for 3 days. D) Comparison of percent of activated receiver mESCs (No Notch, NbALFA-Notch, and NbALFA-SNIPR) when cultured with the different sender mESCs as assessed by fitting a Gaussian mixture model.

In addition to optimizing for rapid iteration, we also sought to introduce a ligand-receptor recognition pair with high-affinity and orthogonality from endogenous cell surface proteins. Previous systems have used GFP or its non-fluorescent mutant nfGFP, which are orthogonal to endogenous proteins but can limit the available channels in fluorescence experiments while also presenting a minor challenge because of the size of the ligand^19,31,32,51–53^. We sought to expand the available orthogonal ligands by using ALFA-tag as our ligand and its cognate binder NbALFA as our antigen sensing domain^54^. We designed NbALFA as the extracellular recognition domain with either the original synNotch or SNIPR (Figure 2A). For our Receiver Cells, we stably integrated synNotch or SNIPR using lentiviral transduction into the same output circuit with TagBFP in mESCs as previously described (Figure 2B). We sorted for similar expression of mCherry expression and surface expression using Myc-tag for NbALFA-synNotch or NbALFA-SNIPR (Supplemental Figure 2A). We designed the ALFA-tag ligand to be displayed on the PDGFR transmembrane domain with a fibcon rigid linker^55,56^ (Figure 2A) downstream of the Ef1a promoter with mCherry as a transduction marker using the Mammalian Cloning Toolkit. We stably integrated this construct into mESCs by lentiviral transduction and sorted a polyclonal cell population using high mCherry expression and surface expression of ALFA-tag (Supplemental Figure 2B). This engineered mESC line (ALFA-mESC) represents the Sender Cell (Figure 2B).

To test for the functionality of our lentivirally generated cell lines, we co-cultured sender and receiver cells at a 1:1 ratio in a low-attachment plate. This culture method allowed homogeneous distribution of our cell types and forced them to grow into well-mixed spheres. We cultured wildtype mESCs (no ligand) alone, ALFA-mESCs alone, output circuit mESCs (No Notch) alone, receiver cells (NbALFA-Notch or NbALFA-SNIPR) alone and set up co-cultures with each of the sender and receiver cells combined. Cultures were assessed after three days for BFP expression by flow cytometry (Figure 2B). We saw that only in co-cultures of ALFA-mESCs with NbALFA-Notch or NbALFA-SNIPR on the receiver cell was there an increase in BFP expression. We saw a robust 1.5-2 log shift in BFP expression in NbALFA-SNIPR versus an approximately half log increase for NBALFA-Notch Receiver Cells (Figure 2C). We quantitatively characterized activation using Gaussian mixture models to separate activated cells from inactive cells in our cytometry histograms. When co-cultured with cells expressing NbALFA-SNIPR, the fraction of cells showing ALFA-tag-dependent BFP expression reached approximately 80% while synNotch was slightly above 40% (Figure 2D). These data indicate that both NbALFA-Notch and NbALFA-SNIPR juxtacrine inducible systems activate BFP expression in mESCs with NbALFA-SNIPR working with greater efficency.

### Engineering GAL4-inducible mESCs to differentiate into glutamatergic neurons

As an application of these systems in the context of cell differentiation, we first sought to engineer mESCs to differentiate into a population of a cell type of interest. Previous work has shown that ectopic expression of the Ngn2 transcription factor is sufficient to drive glutamatergic neuronal differentiation in mESCs^27^. We repeated these studies and verified the same results by transducing mESCs with a plasmid expressing Ngn2 downstream of a constitutive promoter (Figure 3A-B). Differentiation into neuronal cells was confirmed with the neuron-specific class III β tubulin marker using immunofluorescence confocal microscopy.

**Figure 3:**
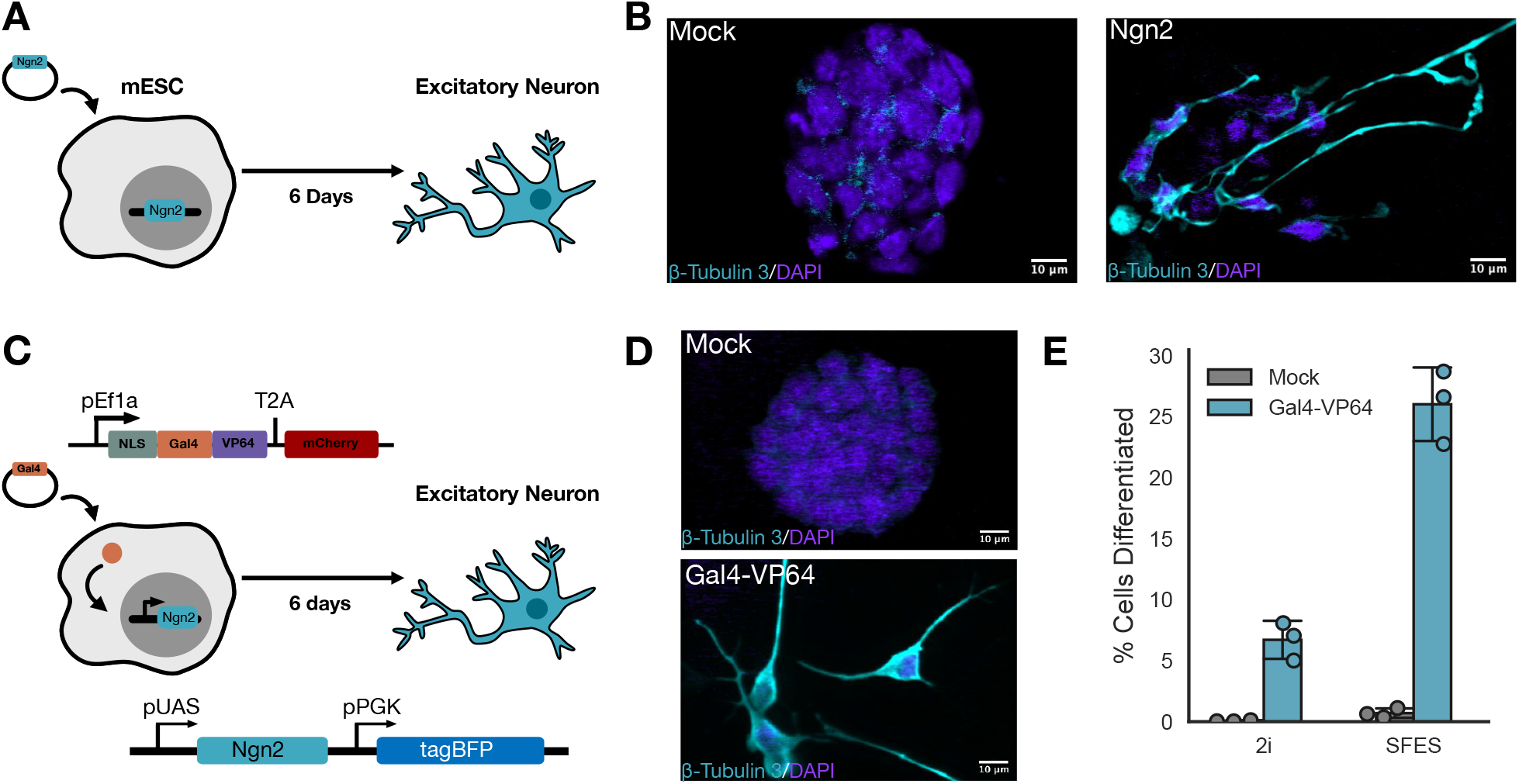
Engineering Gal 4-inducible mESCs to differentiate into glutamatergic neurons. A) Schematic of glutamatergic neuron differentiation program as previously described. B) Excitatory neuron validation using the class III β tubulin marker by confocal microscopy. C) Schematic of glutamatergic neuron differentiation program engineered in mESCs using the transcription factor Ngn2 downstream the inducible UAS promoter. Gal4 transcription factor expressed on a lentiviral expression vector is transduced into the inducible Ngn2 mESC results in expression of Ngn2. D) Excitatory neuron validation using class III β tubulin by confocal microscopy. E) The percent of iNs in either mock transduced or Gal4-VP64 transduced conditions grown in either 2i or SFES media that differentiated as assessed by Gaussian mixture model fitting of population level class III beta tubulin staining.

In order to control differentiation, we engineered mESCs with the Ngn2 transcription factor downstream of an inducible promoter. We expressed Ngn2 downstream of the same GAL4 UAS-ybTATA promoter used above (Figure 3C). We stably integrated this cassette using lentivirus and selected based on constitutive expression of TagBFP as a marker of successful integration into mESCs, producing GAL4-inducible neurons (iNs). We designed a lentiviral expression vector that when transduced into our iNs would express Gal4-VP64, activating Ngn2 expression and differentiation into glutamatergic neurons after six days (Figure 3C). We validated neuronal cells using the neuron-specific class III β tubulin marker with immunofluorescence confocal microscopy (Figure 3D). To determine the efficiency of differentiation in our iNs and exclude the possibility of neurons arising due to random differentiation, we quantified the percentage of class III β tubulin positive cells using flow cytometry in Gal4-VP64 transduced iNs versus mock (viral packaging without Gal4-VP64 plasmid) transduced iNs, as well as Gal4-VP64 transduced wildtype mESCs and mock transduced wildtype mESCs. We cultured in both 2i and SFES media conditions and compared the differentiation efficiency (Figure 3E, Supplemental Figure 3). After Gal4-VP64 transduction, 30% iNs were class III β tubulin positive (Figure 3E, Supplemental Figure 3A-B). Mock transduced iNs (Figure 3E) and wild type mESCs (Supplemental Figure 3C-E) did not differentiate. These data indicate that the GAL4 UAS-ybTATA promoter is capable of driving sufficient expression of Ngn2 to induce neuronal differentiation.

### Orthogonal small molecule synthetic transcription factor systems induce glutamatergic neuronal differentiation in mESCs

We next combined our iNs with each of the small molecule inducible systems. To test for differentiation, we stably integrated each synthetic transcription factor system into separate iN populations by lentiviral transduction (Figure 4A). To validate differentiation, we cultured each system separately for six days at doses roughly equivalent to the EC50 for TagBFP in our initial testing of these systems, using 50nM of 4-OHT, 625nM of ABA and 62.5nM of GZV. We then validated neuronal differentiation with immunofluorescence confocal microscopy (Figure 4B). Neurons were successfully generated in all induced conditions, suggesting that the inducible systems we tested were capable of driving Ngn2-dependent differentiation.

**Figure 4:**
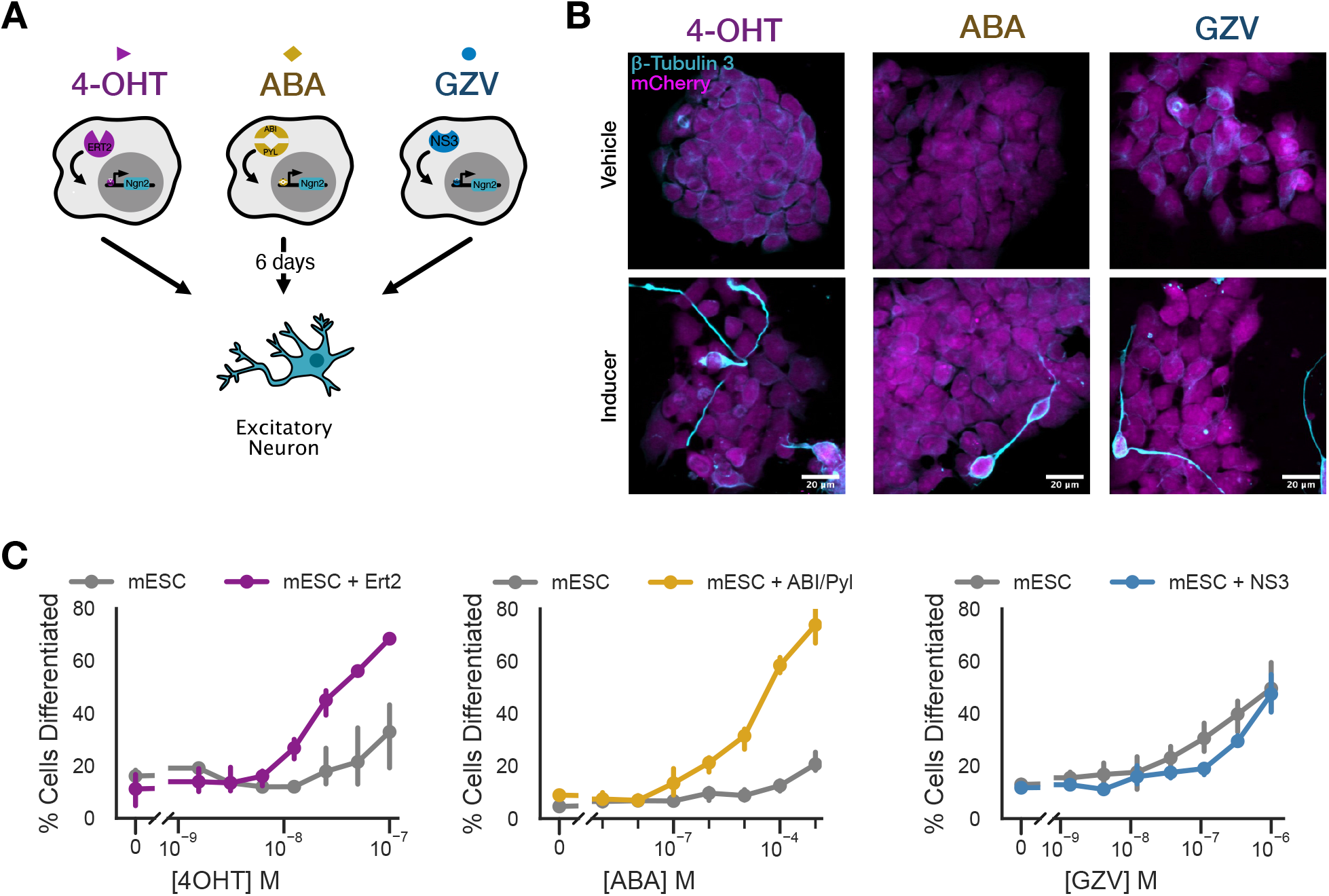
Orthogonal small molecule synthetic transcription factor systems induce glutamatergic neuronal differentiation in mESCs. A) Schematic of assay for small molecule synthetic transcription factor systems with Ngn2 downstream of the inducible system. B) Small molecule-induced differentiation of mESCs into neurons, using 50nM of 4-OHT, 625nM of ABA and 62.5nM of GZV. Validation by staining with the class III β-tubulin marker using confocal microscopy. C) The percent of each synthetic transcription factor iN system that differentiated as assessed by Gaussian mixture model fitting of population level class III beta tubulin staining.

Beyond the role of a master transcription factor, the quantitative requirements of direct differentiation via Ngn2 are poorly understood. We used the dose-dependent expression of our inducible systems to begin to explore the degree to which transcription factor expression level controlled differentiation efficiency. We repeated the above experiment using a range of doses for each inducible system. We quantified differentiation using immunofluorescence staining for class III β-tubulin with flow cytometry (Figure 4C, Supplemental Figure 4A). Both ERT2- and ABI/PYL-inducible systems produced a dose-dependent increase in differentiated cells compared to wild type mESC controls. Notably, 4-OHT treatment induced neuronal differentiation in control cells at high doses, while ABA did not. Curiously, the NS3-inducible system was not able to efficiently generate induced neurons at rates above background. These results indicate that some but not all of these systems are capable of driving direct differentiation of mESCs into neurons with important implications for the quantitative requirements of Ngn2-dependent directed differentiation.

### SNIPR induces glutamatergic neuronal differentiation in mESCs

We sought to test differentiation downstream of juxtacrine induction. Differentiation using synNotch has previously been described^31,32,49^ and we sought to test for differentiation using SNIPR which showed higher expression and percent of cells expressing the receptor’s payload in our hands. We stably integrated NbALFA-SNIPR into our iNs by lentiviral transduction as described above. We then co-cultured the iN or the NbALFA-SNIPR iN receiver cells with wildtype mESC or ALFA-mESC sender cells for six days and validated differentiation by confocal microscopy and flow cytometry (Figure 5A-C, Supplemental Figure 5A-B). We saw differentiation only in co-cultures NbALFA-SNIPR iN receivers and ALFA-mESC senders. Furthermore, in our microscopy assay we observed spatially-dependent patterning where, qualitatively, receivers adjacent to senders seemed more likely to differentiate (Figure 5B). In an effort to increase efficiency of differentiation, we co-cultured at increasing sender to receiver ratios and measured percent differentiation efficiency as a function of density (Supplemental Figure 5B). Increasing sender density did not increase differentiation efficiency, although this effect could be moderated because of the increased number of non-differentiating sender cells in the co-culture. These results indicate that SNIPR is capable of driving direct differentiation of mESCs into neurons in a cell contact- and spatially-dependent manner.

**Figure 5:**
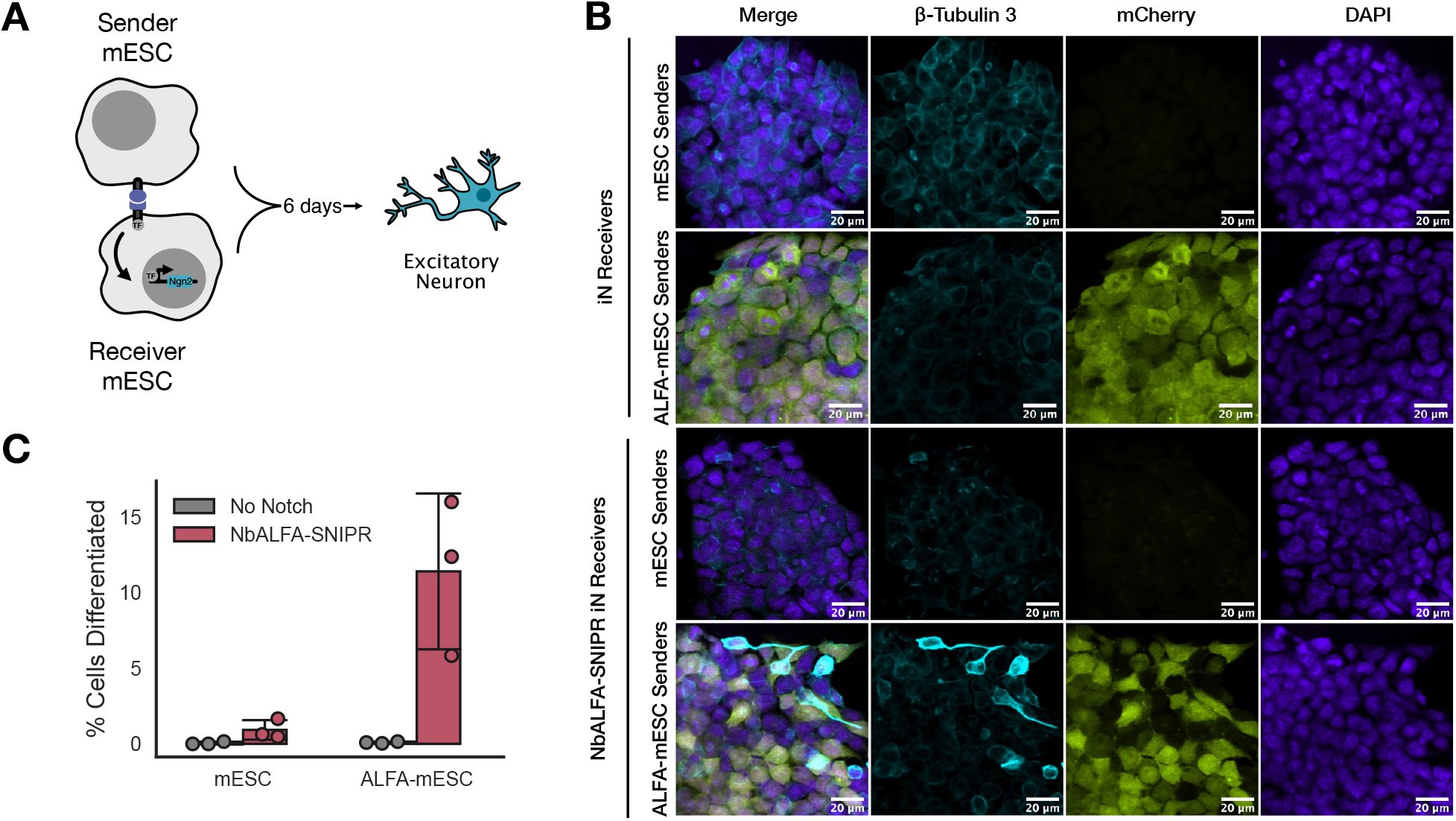
SNIPR induces glutamatergic neuronal differentiation in mESCs. A) Schematic of co-culture assay for ALFA-mESC sender cells and NbALFA-SNIPR receiver cell with Ngn2 downstream of the inducible system (NbALFA-SNIPR iN). B) Differentiation for co-culture assay validated using the class III β-tubulin marker with confocal microscopy. C) The percent of NbALFA-SNIPR iNs when co-cultured with ALFA-mESCs that differentiated as assessed by Gaussian mixture model fitting of population level class III beta tubulin staining.

## Discussion

In this work we demonstrated drug- and cell contact-inducible systems for controlling expression and differentiation in mESCs. We show that three small molecule inducible transcription factor systems as well as two juxtacrine inducible systems work reliably and efficiently in mESCs as measured by expression of BFP output. We demonstrate the functionality of the GAL4 UAS-ybTATA inducible promoter as an orthogonal transcriptional regulator in mESCs. Additionally, we show that when applied in the context of differentiation, some of these inducible systems can drive direct differentiation of mESCs into neurons through driving expression of the transcription factor Ngn2. Through this work, we expand the toolkit for engineering mESCs and provide a rapid, robust, and iterable chassis for engineering mESCs.The tools we present enable quantitative investigation of the requirements for differentiation, and offer multi-dimensional control over development in multiple modalities. Each of the components demonstrated in our study can readily be used on their own or in combination in applications to further our understanding of developmental principles.

Successfully engineering cellular behavior requires considerable iteration, refinement, and optimization. Enabling rapid cycles of cell engineering makes mESCs a more accessible chassis for this work. Our ability to study and understand developmental principles could greatly benefit from tools that expedite the process of asking developmental questions. The lentiviral transduction system we report here, compatible with the previously published Mammalian Toolkit for rapidly generating genetic components for cell engineering, enables rapid, robust, and iterable engineering workflows in mESCs This system complements high efficiency landing pads^21,57,58^ for cases where controlling copy number is essential.

The different abilities of inducible systems in inducing differentiation begs questions about quantitative requirements for differentiation. For example, what is the minimal and maximal amount of a reprogramming transcription factor needed to be in the “ON” state for differentiation to occur? How do the kinetics of transcription factor production contribute to differentiation? How do transcriptional and proteostatic dynamics contribute to the decision to differentiate? The different mechanisms of the inducible systems used here might suggest an answer, whereby the only system that requires de novo synthesis of its transcription factor was also the only system unable to stimulate differentiation. Future work should explore differences in overall payload production between these inducible systems. Further, the off-target differentiation seen in wild type cells treated with 4-OHT raises questions about this drug’s orthogonality and suggest dosing regimes to minimize its off-target effects in synthetic developmental studies.

The tools we present in this manuscript are essential for enabling the next generation of synthetic developmental biology. Because of the complexity of development, synthetic development requires multi-dimensional control over multiple scales necessary to understand and build more complex developmental systems. The tools presented here add to a growing toolkit that enables this multidimensional multiscale control over cellular behavior. Although we show the efficacy of these tools in a relatively simple expression system to direct mESC into neurons, these tools are ready to be integrated into more complex synthetic circuitry such as feedback controllers^59,60^. Additionally, implementing these systems into building more controlled and refined in vitro culture models like organoids by controlling cell fate, timing of differentiation, and organization, will give us a deeper understanding of tissue composition and ultimately of predictive tissue engineering^8^.

## Materials and Methods

### Cell Culture

mESCs were cultured in serum free ES media (SFES) supplemented with 2i and LIF as described in Mulas et al 2019^61^. SFES media consists of 500 mL DMEM/F12 (Thermo #21331020), 500 mL Neurobasal (Gibco #21103-049), 5 mL N2 Supplement (Gibco #17502-048), 10 mL B27 (Thermo #17504044), 6.66 mL 7.5% BSA (Gibco #15260-037), 10 mL 100x GlutaMax (Gibco #35050-061), 1mL 55mM 2-Mercaptoethanol and 10mL of Antibiotic-Antimycotic. To make “2i” medium, 1 uM PD03259010 (Selleckchem #S1036), 3 uM CHIR99021 (Selleckchem #S2924) and 10^4 units/mL LIF (ESGRO #ESG1107) were added to 45 mL SFES. To passage, mESCs were treated with 0.5 mL of accutase in a 6 well plate (Corning #353046) for 5 minutes at room temperature. After incubation, cells were mixed by pipette and moved to a 15 mL conical tube, supplemented with 10 mL wash media and spun at 400xg for 4 minutes. Wash media consists of DMEM/F12 (Thermo #21331020) supplemented with 8mL 7.5% BSA. Then, wash media was removed and cells were counted using the Countess II Cell Counter (ThermoFisher) according to the manufacturer’s instructions. Cells were then plated in 6 well plates that had gelatinized with 1% gelatin for 30 minutes at 37°C at 1.5e5 cells per well in 2 mL of 2i. Cells were split every other day or three days (1e5 cells). All cells were maintained at 37°C with 5% CO2.

Lenti-X 293T packaging cells (Clontech #11131D) were cultured in medium consisting of Dulbecco’s Modified Eagle Medium (DMEM) (Gibco #10569-010) and 10% fetal bovine serum (FBS) (University of California, San Francisco [UCSF] Cell Culture Facility). Lenti-X 293T cells were cultured in T150 or T225 flasks (Corning #430825 and #431082) and passaged upon reaching 80% confluency. To passage, cells were treated with TrypLE express (Gibco #12605010) at 37°C for 5 minutes. Then, 10 mL of media was used to quench the reaction and cells were collected into a 50 mL conical tube and pelleted by centrifugation (400xg for 4 minutes). Cells were cultured until passage 30 whereupon fresh Lenti-X 293 T cells were thawed. All cells were maintained at 37°C with 5% CO2.

### DNA Constructs

The output circuit, pHR_Gal4UASpyb‐TATA_tBFP_pGK_mCitrine and the SNIPR plasmid were generous gifts by Dr. Kole Roybal. The small molecule synthetic transcription factors and the sender and receiver juxtacrine plasmids were constructed using the Mammalian Toolkit (MTK)^21^, a hierarchical DNA assembly method based on Golden-Gate (GG) cloning^62^. The SNIPR used in this study contains the CD8a hinge domain, the human Notch1 transmembrane domain, and the human Notch2 juxtamembrane domain. The SNIPR was domesticated as an MTK part 3B with the Gal4-VP64 transcriptional actuator domain. The juxtacrine sender cell construct was assembled using MTK by domesticating ALFA epitope tag^54^ to a single fibcon^56^ domain as an extracellular scaffold as a part 3A and the domesticating the PDGFRb transmembrane domain as a part 3B. The receiver cell plasmid consisted of the standard SNIPR core with an anti-ALFA tag nanobody as a binder. All constructs were assembled into a novel lentiviral destination backbone^50,55^ via a PaqCI reaction following manufacturer instructions (New England Biolabs #R0745S). The Gal4-NGN2 plasmid was constructed using Gibson Assembly with In-Fusion® Snap Assembly Master Mix (Takara Bio #638947) and following recommended manufacturer’s instructions. All plasmids were propagated in Stbl3 E. coli (QB3 MacroLab). Domestication was verified via sequencing and transcriptional unit assembly was verified via restriction digest. All plasmids will be available on Addgene. The sequence for key constructs used in this study are included as supplemental data and described in Supplemental Table 1.

### Lentiviral Transduction and Generation of Stable Cell Lines

Lenti-X 293T cells (Takara Bio #632180) were seeded at approximately 7e5 cells/well in a 6-well plate to yield ∼80% confluency the following day. The following day cells were transfected with 1.5μg of transfer vector containing the desired expression cassette, and the lentiviral packaging plasmids pMD2.G (170ng) and pCMV-dR8.91 (1.33μg) using 10ul of Fugene HD (Promega #E2312), according to manufacturer procedure. At 48 hours the viral supernatant was filtered through a 0.45μm PVDF filter and concentrated using the Lenti-X concentrator (Takara, #631231) according to the manufacturer’s instructions. Lenti-X concentrator solution was added at a 1:3 viral supernatant:concentrator ratio, mixed by inversion, and incubated at 4 C for at least 2 hours. Supernatant-concentrator mix was pelleted by centrifugation at 1500xg at 4 C for 45 minutes, supernatant was removed and pellet was resuspended using 100 uL 2i media for each T25 flask of mESCs. 2 wells of a 6 well plate was concentrated and applied to 1 T25 flask plated with 500K cells on the day of transduction. 72 hours after the addition of the viral supernatant to mESC culture, polyclonal cell populations were selected via fluorescence-activated cell sorting (FACS) as described below. Cells were sorted using BD FACS Aria sorter.

### Small Molecule Inducible System Assays (BFP)

Stock solutions of abscisic acid (Sigma Aldrich #862169, 50 mM in ethanol), 4-hydroxytamoxifen (Sigma Aldrich #H6278, 1 mM in ethanol), and grazoprevir (MedChemExpress #HY-15298, 1 mM in DMSO) were stored at -80C. 1e4 mESCs were seeded into 96-well gelatin coated flat-bottom plates in 100 uL 2i media on day 0 of induction. On day 1 of induction,100 uL of media containing 2X concentrated amounts of 4-OHT, 10X of ABA, and 3X of GZV was added to maximum dosage row of wells and serial dilution was performed across each row, ending on only diluent on minimum dosage row. On day 4, cells were lifted and transferred to 96 well round bottom plates and prepared for flow cytometry as described below. All inductions were performed in biological triplicates.

### Small Molecule Inducible System Assays (NGN2)

500 mESCs were seeded into 96-well gelatin coated flat-bottom plates in 100 uL 2i media on day 0 of induction. On day 1 of induction, 2i media was exchanged for 100 uL of SFES media. 100uL of SFES containing 2X concentrated amounts of 4-OHT, 10X of ABA, and 3X of GZV was added to maximum dosage row of wells and serial dilution was performed across each row, ending on only diluent on minimum dosage row. On Day 4, media was refreshed with fresh induction media. On day 6, cells were prepared for intracellular staining and flow cytometry as described below. All inductions were performed in biological triplicates.

### Antibody Staining Live Cells

All experiments using antibody staining were performed in 96 well round bottom plates. After dissociation, cells for these assays were pelleted by centrifugation (400xg for 4 minutes) and supernatant was removed. Cells were resuspended for 45 minutes with appropriate antibodies in a staining solution of 25 uL PBS. After incubation, plates were spun down, rinsed with DPBS, and resuspended in flow buffer made of DPBS with 2% FBS and 2mM EDTA (Thermo #AM9260G).

### Antibodies

Antibodies used for live cell flow cytometry assays include Alexa Fluor 647-Anti-Myc tag conjugated antibody (Cell Signaling Technologies #2233S) used at 1:100, FluoTag®-X2 anti-ALFA Atto-488 conjugated antibody used at 1:50. All antibodies were diluted in DPBS (UCSF Cell Culture Facility) for staining. For FACS, all antibodies were used at 1:50 in 200 uL.

Antibodies for fixed intracellular staining assays include FluoTag®-X2 anti-ALFA Atto-488 (NanoTag Biotechnologies #N1502-At488-L) conjugated antibody diluted at 1:50 and Anti-beta III Tubulin antibody [EP1569Y]-Alexa Fluor 647 (ab190575) conjugated antibody used at 1:500. All antibodies were diluted in blocking buffer as described in the Bioscience Foxp3/Transcription Factor Staining Buffer Set (ThermoFisher #00-5523-00).

Antibodies used for immunofluorescence include anti-beta III Tubulin antibody [EP1569Y] (Abcam ab52623) used at 1:500 dilution, Alexa Fluor 647 secondary antibody (Thermo Scientific #A-21244) used at 1:2000. All antibodies were diluted in blocking buffer.

### FACS

Cell lines were bulk sorted for high expression using the UCSF Laboratory for Cell Analysis Core Facility FACSAriaII (BD Biosciences). Cells were assessed for BFP (405nm excitation, 525/50nm emission, 505lp collection dichroic), mCitrine (488nm ex, 530/30nm em, 505lp cd), mCherry (561nm ex, 610/20nm em, 600lp cd), and Alexa647 (633nm ex, 670/30nm em) fluorescence. Fluorescence-negative controls were used to set detector power so that negative cells appeared to have mean fluorescence ∼100 counts, and then transduced cells were sorted for cells with expression outside of the negative control expression level. All data were collected using FACSDiva (BD Biosciences).

### Flow Cytometry

All flow cytometry data was obtained using a LSR Fortessa or LSRII (BD Biosciences). All assays were run in a 96-well round bottom plate (Fisher Scientific #08-772-2C). Samples were prepared by pelleting cells in the plate using centrifugation at 400xg for 4 minutes. Supernatant was then removed and 200 uL of PBS (UCSF Cell Culture facility) was used to wash cells. The cells were again pelleted as described above and supernatant was removed. Cells were resuspended in 200 uL of flow buffer (DPBS with 2% FBS and 2mM EDTA) and mixed by pipetting prior to flow cytometry assay. At least 1e7 events were recorded for all experiments.

### synNotch and SNIPR activation assays

mESCs expressing synNotch and SNIPR constructs were seeded at a density of 5e4 cells/well in a low-attachment round bottom 96-well plate (Corning #7007). Cells were plated alone or with an equal number of either WT mESCs or mESCs expressing the ALFA-tag ligand in a total of 200μl of 2i medium. Plates were spun briefly (400xg for 1 minute) to increase likelihood of cell-cell interaction. Cells were co-cultured for 72 hours and the BFP expression and surface expression of receptors and ALFA-tag were conducted using flow cytometry. synNotch and SNIPR activation was assessed as described previously by fitting a two-component Gaussian mixture model to BFP expression data and estimating the fraction of the population in the ‘off’ and ‘on’ components^41^.

### SNIPR Activation-Ngn2 Differentiation Assay

mESCs expressing SNIPR constructs were seeded at a density of 1e4 cells/well in a low-attachment round bottom 96-well plate. Cells were plated alone or at 1:1, 2:1, and 4:1 ratios with mESCs expressing the ALFA-tag ligand in a total of 200μl of 2i medium. Plates were spun briefly (400xg for 1 minute) to increase likelihood of cell-cell interaction. Cells were co-cultured for 6 days then fixed as described above. Beta III Tubulin expression and surface expression of receptors and ALFA-tag were conducted using flow cytometry. SNIPR activation and differentiation was assessed as described previously by fitting a two-component Gaussian mixture model.

### Ngn2-dependent direct differentiation

Gal4-Ngn2 iNs and wildtype cells were seeded at a density of 100k in 6-well plate of 2i media the day before viral infection. On the day of transduction, one well of a 6-well plate of virus containing the Gal4 plasmid was concentrated as described above and added to one well of Ngn2 and wildtype cells. Mock transduced iNs and wildtype cells underwent the same procedure. Cells were incubated for 48 hours then polyclonal populations were sorted for expression of TagBFP and mCherry. Cells were then plated in gelatin coated 96-well flat bottom plate at 1e3 cells/well in 2i medium. The following day media was refreshed in cells under 2i media conditions and exchanged with SFES in the case of SFES media conditions. Cells were incubated for at least six days with media refreshed every two days. Cells were then fixed and analyzed by flow cytometry as described above.

### Immunofluorescence and fixed cell microscopy

Glass coverslips (Fisher Scientific #12-545-81P) were coated with 5ug/ml Fibronectin (Sigma-Aldrich #F0895-1MG) in HBSS (Fisher Scientific #14-025-092) and placed in 24 well plate (Fisherbrand #FB012929). Cells were lifted from 96 well plates and were placed with 2i media and cultured for an additional 3 days. Cells were fixed in 4% PFA/PBS for 20 minutes. Blocking and permeabilization were performed in a blocking solution consisting of 0.2M glycine, 2.5% FBS, 0.1% Triton X-100 in 1X PBS for 30 minutes at room temperature. Primary and secondary antibodies were diluted and incubated in the blocking solution. Cells were incubated with primary antibodies for 30 minutes at room temperature and washed 3x with PBS. Cells were then incubated with secondary antibodies for 30 minutes at room temperature and washed 3x with PBS. Coverslips were then mounted in ProLong DAPI Fluoromount (Thermo Scientific #P36966) onto slide. Images were collected using Nikon Ti inverted microscope equipped with Yokogawa CSU-22 spinning disk confocal and a custom 4-line solid state laser launch (100 mW at 405, 488, 561, and 640 nm excitation) using 40x air objective and processed using ImageJ/Fiji.

### Fixed Cell Staining to Assess Differentiation

Media was removed from cells in 96-well flat bottom plate. Cells were treated with Zombie UV Viability Kit (Biolegend #280.423107) diluted 1:500 in PBS for 30 minutes in the dark at room temperature. Cells were lifted with accutase/mESC wash and transferred to a 96-well round bottom plate. Cells were pelleted at 400xg for 4 minutes and supernatant was removed. Cells then underwent fixation and permeabilization using an Biosciences Foxp3/Transcription Factor Staining Buffer Set (ThermoFisher #00-5523-00) following manufacturer’s recommended procedure. Following the fixation/permeabilization process, cells were stained for 1 hour at room temperature in the dark. The stain was then washed off with the perm/wash buffer in the Biosciences kit twice and resuspended in flow buffer. Cells were analyzed by flow cytometry.

### Data Presentation, Analysis, and Availability

All experiments were performed in at least biological triplicate. Central tendency for individual replicates are presented as circles where possible. For BFP expression experiments, raw fluorescence is reported as median fluorescence value for all cells within a single replicate, and the presented means and error are calculated between replicates. For class III beta tubulin staining experiments, raw fluorescence is reported as mean fluorescence value for all cells within a single replicate and the presented means and error are calculated between replicates. For live cell flow cytometry, collected events were filtered to remove small events and then gated on FSC and SSC to capture singlet populations. For fixed cell flow cytometry, small events were filtered out. Histogram density estimates represent all events for all 3 replicates in a flow cytometry experiment, and are calculated via seaborn’s kdeplot function with bandwidth adjustment of 0.2. All data analysis was conducted using custom Python scripts, available on github^63^. Analysis was conducted in Jupyter^64^ and relied on numpy^65^, matplotlib^66,67^, seaborn^68^, pandas^69,70^, SciPy^71^, scikit-learn^72^ and fcsparser. ALFA tag and NbALFA schematics were inspired by their published structures and presented using elements from a proposed protein emoticon^73^. All primary data from flow cytometry experiments and immunofluorescence microscopy are available on Zenodo^74^. All figures assembled with Affinity Designer 2.

## Supporting information

Supplemental DNA Sequences

Supplemental Table 1 - DNA Constructs

## Acknowledgments

The authors would like to thank Dr. Wendell A. Lim, Dr. Todd G. Nystul, and all members of the El-Samad and Lim labs for helpful and essential discussion. Data for this study were acquired at the Laboratory for Cell Analysis and Center for Advanced Light Microscopy at UCSF. ZYW thanks Bully and Stephanie E. Crilly for essential support and advice.

## Author Contributions

SSS: Conceptualization, investigation, methodology, formal analysis, resources, writing - original draft, writing - review & editing. DHS: resources, writing - review & editing. HE-S: Conceptualization, supervision, resources, funding acquisition, writing - review & editing. ZYW: Conceptualization, supervision, methodology, resources, formal analysis, funding acquisition, writing - review & editing.

