## Supplemental Figures for "Small molecule and cell contact-inducible systems for controlling expression and differentiation in stem cells"

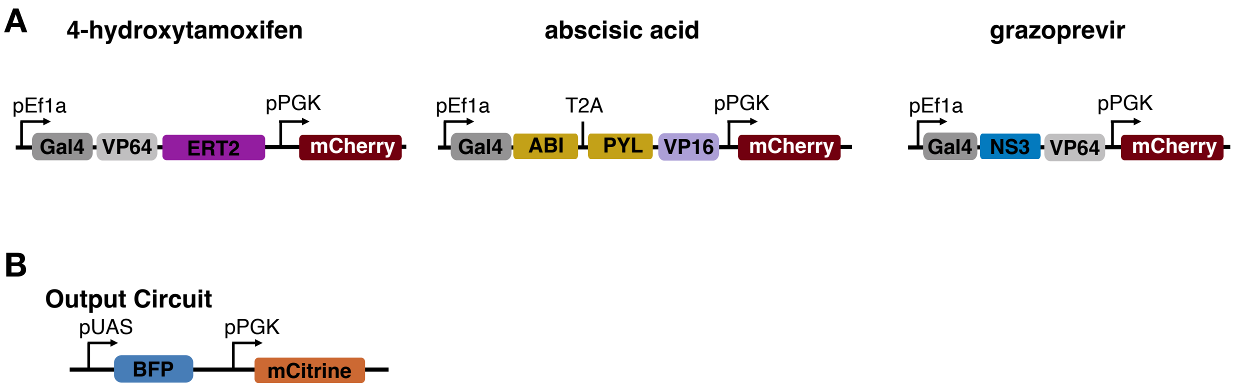


**Supplemental Figure 1: Schematic Representation of Constructs.** A) Schematic of each drug-inducible construct with a constitutively expressed mCherry. B) The output circuit, with TagBFP expression downstream of the (5X)UAS ybTATA promoter and a constitutively expressed mCitrine.

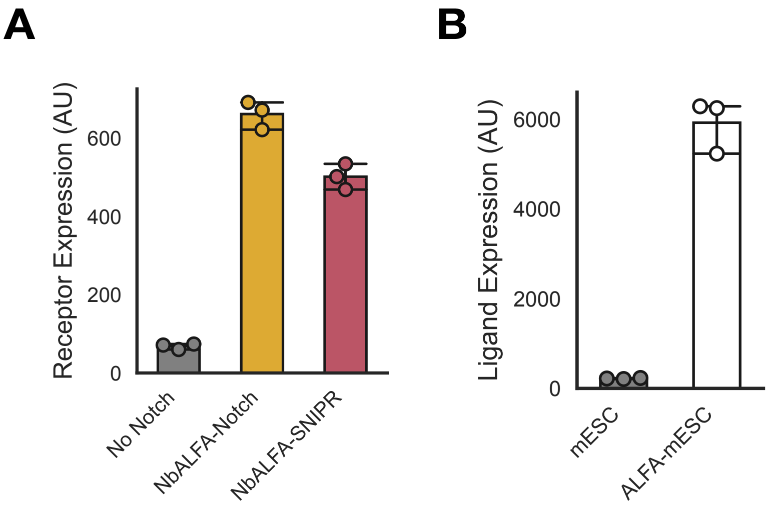


**Supplemental Figure 2: Juxtacrine Receptor and Ligand Expression Levels.** A) Comparison of median receptor expression levels across output circuit mESCs (No Notch), synNotch (NbALFA-Notch), and SNIPR (NbALFA-SNIPR)-expressing mESCs as assessed by surface expression using anti-Myc staining. B) Comparison of median ligand expression levels between wildtype mESCs and ALFA-mESCs as assessed by surface expression of ALFA-tag using anti-ALFA-tag staining.

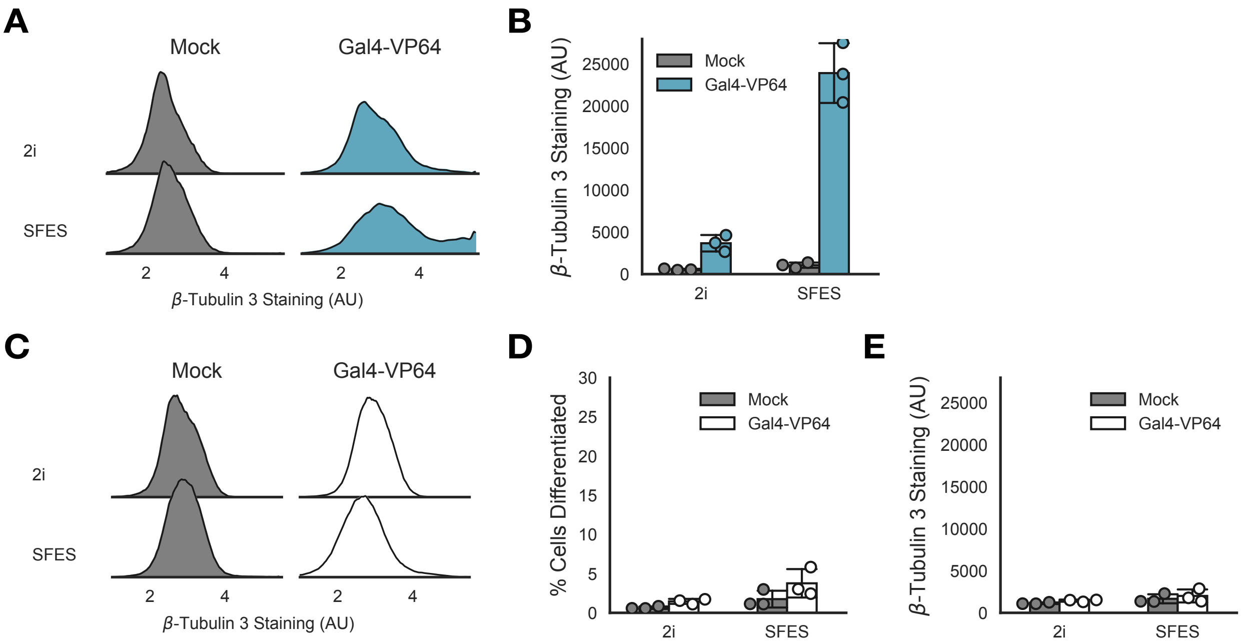


**Supplemental Figure 3: Quantification of iNs.** A) Population density plots comparing class III beta tubulin staining in iNs in either mock transduced or Gal4-VP64 transduced conditions grown in either 2i or SFES media. B) Mean class III beta tubulin staining in iNs mock transduced or transduced with Gal4-VP64 and cultured in either 2i or SFES. C) Population density plots comparing class III beta tubulin staining in wildtype mESCs in either mock transduced or Gal4-VP64 transduced conditions grown in either 2i or SFES media. D) The percent of wildtype mESCs that differentiated as assessed by Gaussian mixture model fitting of population level class III beta tubulin staining. E) Mean class III beta tubulin staining in wildtype mESCs.

**A**

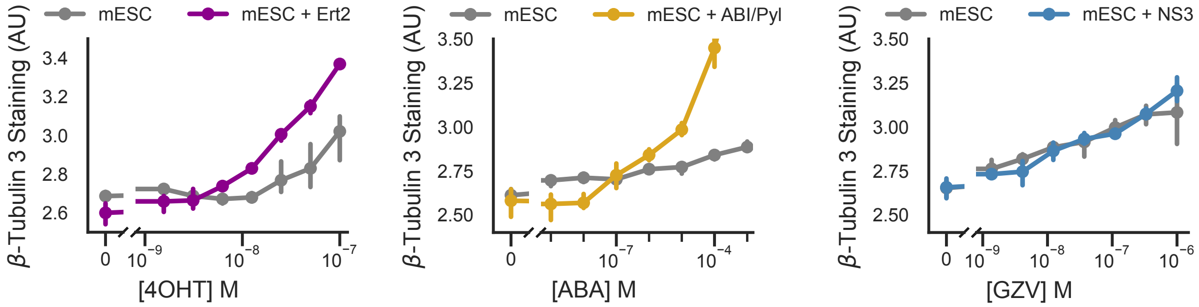


**Supplemental Figure 4: Class III β-tubulin staining for small molecule driven iNs.** A) Median class III β-tubulin staining for each inducible driven iN system as measured by flow cytometry.

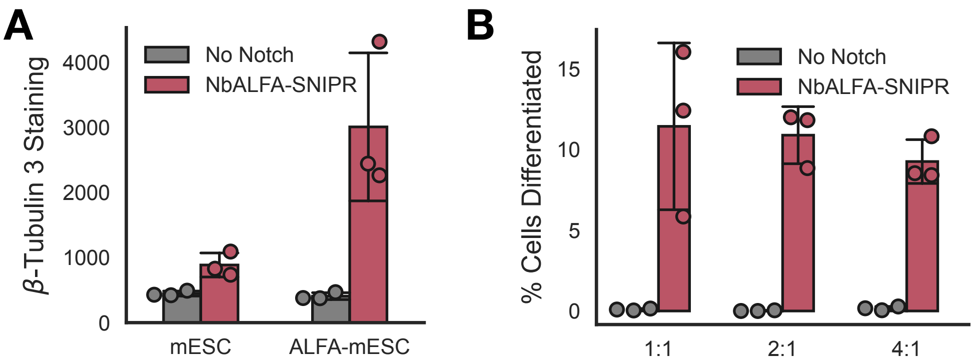


**Supplemental Figure 5: Quantification of NbALFA-SNIPR iNs.** A) Median class III β-tubulin staining in NbALFA-SNIPR iN differentiation assay as measured by flow cytometry. B) The percent of NbALFA-SNIPR iNs that differentiated as assessed by Gaussian mixture model fitting of population level class III beta tubulin staining across increasing sender cell ratios and as measured by class III β-tubulin and flow cytometry.
